# Dendritic varicosities revealed as important microtubule organisers in neurons

**DOI:** 10.64898/2026.07.13.738214

**Authors:** Adrià Chorro, Chithran Vineethakumari, Paul T. Conduit

## Abstract

Microtubules are polarised polymers that assemble into highly specialised networks in a cell-specific manner. This is controlled in part by microtubule organising centres (MTOCs), which concentrate factors necessary for microtubule nucleation and the organisation of microtubule minus ends. Neurons rely on oppositely polarised microtubule networks, with axons containing mostly plus-end-out microtubules, and dendrites contain many minus-end-out microtubules. How minus-end-out microtubule polarity is established in dendrites remains an important question. Here, we identify a new type of MTOC within the dendrites of *Drosophila* class I dendritic arborisation neurons, a common model for the neuronal cytoskeleton. We show that membrane swellings distributed intermittently along dendrite shafts, which we term “dendritic varicosities”, contain the principal component of the microtubule nucleating complex and repeatedly generate microtubules whose plus ends grow back towards the soma. Varicosities located specifically in distal regions also contain MTOC proteins implicated in minus end anchoring, and this correlates with the accumulation of minus ends specifically in distal varicosities. Depletion of these MTOC proteins leads to major defects in minus end organisation, with microtubule buckles and loops deforming the neuronal membrane. Thus, dendritic varicosities are an important new type of neuronal MTOC that contribute to the generation and organisation of the minus-end-out microtubule network within dendrites.

## Introduction

Microtubules are polarised polymers with a dynamic plus end are a typically more stable or capped minus end. Minus ends are often embedded in microtubule organising centres (MTOCs), which recruit and concentrate proteins required for microtubule nucleation, polymerisation, and anchoring (Sanchez and Feldman, 2016). The best characterised MTOC is the mitotic centrosome (Conduit et al., 2015), but non-centrosomal MTOCs (nc-MTOCs), such as the surfaces of Golgi, endosomes, mitochondria and the nuclear envelope, frequently contribute to microtubule organisation in differentiated cells (Akhmanova and Kapitein, 2022). Different cells employ different MTOCs to organise their highly specialised microtubule networks (Akhmanova and Kapitein, 2022; Sallee and Feldman, 2021), and identifying these MTOCs and how they are regulated remains an important question in cell biology.

A primary component of MTOCs are multi-protein γ-tubulin ring complexes (γ-TuRCs), which template and stimulate microtubule nucleation (Tovey and Conduit, 2018). During nucleation, the minus end remains associated with the γ-TuRC, while the plus end grows out and away, providing a means to control microtubule polarity within cells. Minus ends may remain associated with γ-TuRCs and thus the MTOC, but they can also be released to either grow, shrink, or become stabilised by other proteins (Vineethakumari and Lüders, 2022). Experiments in vitro have shown that minus end release can occur either by microtubule severing (Henkin et al., 2023) or by competition from CAMSAP proteins (Patronin in *Drosophila*) (Rai et al., 2024). Patronin/CAMSAP proteins counteract the depolymerising activity of atypical Kinesins, allowing slow minus end growth (Goodwin and Vale, 2010; Wang et al., 2013; Hendershott and Vale, 2014; Jiang et al., 2014; Feng et al., 2019) or anchoring (Wu et al., 2016; Nashchekin et al., 2016). Minus end anchoring can also be regulated by Ninein proteins (Vineethakumari and Lüders, 2022), which can bind γ-TuRCs, microtubules and function as an adaptor for the minus end directed motor Dynein (Casenghi et al., 2005; Kowanda et al., 2016; Zheng et al., 2016; Redwine et al., 2017).

Neurons are particularly interesting as they rely on long, decentralised and highly polarised microtubule networks that facilitate neuronal growth, branching and stability (Kapitein and Hoogenraad, 2015). An essential feature is that nearly all microtubules within axons have their plus ends pointing away from the soma (plus-end-out), while a large proportion of microtubules in dendrites have their minus ends pointing away from the soma (minus-end-out) (Kapitein and Hoogenraad, 2015). This distinct polarity is ‘read’ by molecular motors, ensuring cargo is delivered to the appropriate compartment to maintain axonal and dendritic identity (Nirschl et al., 2017; Tas et al., 2017; Thyagarajan et al., 2022). While plus-end-out microtubule polarity in axons can be established simply by plus end growth from an MTOC within the soma, establishing minus-end-out polarity in dendrites is less trivial. One non-exclusive mechanism is through local nucleation within dendrites, where the plus ends of newly formed microtubules can naturally grow back towards the soma. Which type of nc-MTOCs control this remains an open question.

*Drosophila* class I dorsal dendritic arborisation E (ddaE) neurons are a common model for studying the neuronal cytoskeleton and microtubule nucleation has been proposed to occur via endosome-associated γ-TuRCs within dendritic branchpoints (Weiner et al., 2020). These branchpoints, however, are far from the distal terminal ends of the long secondary dendrites of these “comb-shaped” arbors, suggesting that other, more distal, nc-MTOCs may exist. Here, we report that local expansions of membrane (varicosities) distributed along the secondary dendrites function as nc-MTOCs. A certain fraction of varicosities contain γ-tubulin and nucleate microtubules that grow back towards the soma. Minus ends accumulate specifically in distal varicosities, where the minus end organising proteins Cnn and Ninein also accumulate. Co-depletion of Cnn and Ninein results in microtubule buckling and looping within expanded varicosities close to the dendrite tips, along with defects in overall microtubule polarity. Collectively, the data reveal that dendritic varicosities represent a new type of nc-MTOC that helps regulate the minus-end-out microtubule network in dendrites.

## Results and Discussion

### Dendritic varicosities are sites of microtubule nucleation

Proprioceptive larval *Drosophila* class I ddaE neurons have “E-shaped” dendritic arbors (Fig. 1A) that use mechanosensory ion channels to sense stretching and compression during larval crawling (Guo et al., 2016; He et al., 2019). We assessed microtubule dynamics in the relatively straight secondary dendrites, which have an average length of ∼107mm, ranging from ∼33mm to ∼188mm (n = 108 dendrites) (Fig. 1A). Expressing End-binding protein 1-GFP (EB1-GFP) in these neurons labels both growing plus and minus ends, producing bright fast-moving (>2.9mm/min) plus end ‘comets’ and dim slow-moving minus end ‘comets’ (Feng et al., 2019) (Fig. 1B,C). The speed of comets correlated well with the distance of their growth – fast plus ends grew an average of 5.8mm with a maximum distance of 46mm, while slow minus ends grew on average only 2.9mm with a maximum distance of only 7.8mm (Fig. 1D). Dendrites displayed predominantly minus-end-out polarity, with the vast majority of plus and minus ends in proximal and medial regions growing retrograde and anterograde, respectively (Fig. 1E). Nevertheless, even in these “correctly polarised” dendrites, microtubules grew in both directions with roughly equal frequency in distal regions (Fig. 1E).

**Figure 1.**
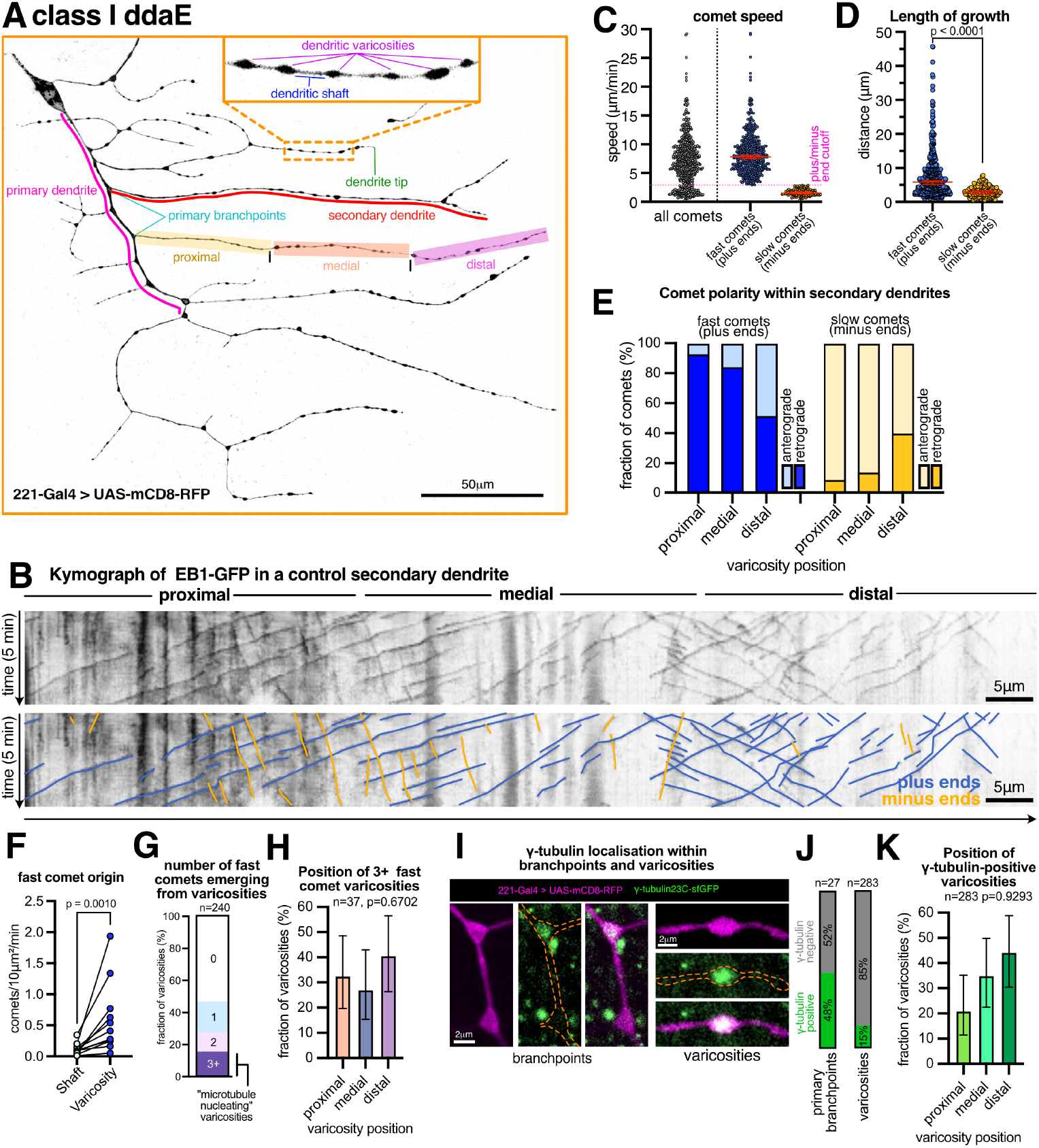
Dendritic varicosities are sites of microtubule nucleation. (**A**) A dendritic arbor of a living 3rd instar larval class I ddaE neuron expressing the membrane marker UAS-mCD8-RFP (inverted gray-scale) with 221-Gal4. Certain features are indicated: primary and secondary dendrites, primary branchpoints, and the dendrite tip. An inset shows a selection of dendritic varicosities. (**B**) An example kymograph from a 5-minute video of a secondary dendrite of a neuron within a living 3rd instar larva expressing UAS-EB1-GFP with 221-Gal4. The bottom panel is a duplicate of the top panel with blue and yellow lines indicating fast plus end and slow minus end growth, respectively. (**C**) Graph displays the speed of individual EB1-GFP comets within secondary dendrites. A speed cutoff of 2.9µm/min was set based on the two apparent distributions and the fast plus and slow minus end comets were re-plotted separately (medians with 95% CI are shown). (**D**) Graph displaying the length of fast (plus ends) and slow (minus ends) EB1-GFP comets in secondary dendrites. Medians with 95% CIs are shown. A lognormal Welch’s t-test was used to compare the data. (**E**) Graph displays the proportion of fast (blue) or slow (yellow) EB1-GFP comets that emerge within proximal (0-33%), medial (34-66%) or distal (67-100%) sections of the secondary dendrite and grew either retrograde (dark tone) or anterograde (light tone). (**F**) Graph displays the frequency at which fast EB1-GFP comets emerged from either shafts (light tone) or varicosities (dark tone). Each point represents the value from an individual dendrite. Lines join data from the same dendrite. Wilcoxon matched-pairs signed rank tests were used to compare the data from shafts and varicosities. For graphs (C) to (F) data obtained from 479 fast and 129 slow comets within 10 secondary dendrites from 8 neurons. (**G**) Graph showing the proportion of varicosities classified based on the number of EB1-GFP comets they produced. (**H**) Graph displaying the spatial distribution of varicosities that have 3 or more EB1-GFP comets emerging from them. A Chi^2^ test was performed to compare the observed proportions to what would be expected based on the total number of varicosities in each section. (**I**) Confocal images show branchpoints and varicosities from living 3^rd^ instar larval class I ddaE neurons expressing the membrane marker UAS-mCD8-RFP (magenta) with 221-Gal4 along with endogenously tagged γ-tubulin-sfGFP (green). Note how accumulations of γ-tubulin-sfGFP signal can be observed within branchpoints and varicosities. The green foci outside of the neuron represent either γ-tubulin accumulations in neighbouring epithelial cells or autofluorescence. (**J**) Graphs displaying the proportion of primary branchpoints or varicosities containing a γ-tubulin-sfGFP signal above cytosolic background. (**K**) The proportion of varicosities that contain a γ-tubulin-sfGFP signal >20% above cytosolic background levels in relation to varicosity position. N=283 varicosities from 23 dendrites from 3 neurons. A Chi^2^ test was performed as in (H). Note how there is no correlation, showing that there is an equal probability of γ-tubulin-sfGFP localising in proximal, medial and distal varicosities. Related to Figure S1.

We had previously reported the presence of local membrane swellings – that we term dendritic varicosities – intermittently distributed along the length of these secondary dendrites (inset, Fig. 1A) (Mukherjee et al., 2020). Structurally, these sites resemble varicosities in mammalian axons and radial glia, both of which serve as nc-MTOCs (Qu et al., 2019; Coquand et al., 2021). Visually identifiable varicosities had a diameter at least 1.7-fold larger than the neighbouring shaft. They appeared on average every ∼8.5mm, with their numbers increasing roughly linearly with dendrite length (Fig. S1A), being more common in distal regions (Fig. S1B) and with one in every three dendrite tips displaying a varicosity (Fig. S1C). These varicosities are observed in both live and fixed samples, including after rapid heat fixation (Fig. S1D), which can better preserve tissue structure (Chen and Johnston, 2022).

Correlating the origin of fast moving EB1-GFP comets with the position of varicosities revealed a strong preference for plus ends to initiate growth within varicosities compared to dendritic shafts that lie between varicosities (Fig. 1F; Video S1). Moreover, ∼15% of varicosities had at least three EB1-GFP comets emerging during the course of imaging (Fig. 1G). Repeated comet emergence is a hallmark of microtubule nucleation and something unlikely to occur due to stochastic regrowth. We considered these ‘3+ fast comet’ varicosities as ‘nucleating’ varicosities and found they were located in either proximal, medial or distal regions (Fig. 1H). Even within individual dendrites, nucleating varicosities could be positioned in multiple different regions (Fig. S1E). Comets emerging from either proximal or medial nucleating varicosities showed a strong retrograde bias, travelling predominantly towards the soma, while comets emerging from distal nucleating varicosities moved either retrograde or anterograde with roughly equal frequency (Fig. S1F). The reason for this difference remains unclear.

Consistent with ∼15% of varicosities being sites of microtubule nucleation, ∼15% contained endogenously-tagged γ-tubulin-GFP signal above cytosolic background (Fig. 1I,J), with γ-tubulin-positive varicosities also located in either proximal, medial or distal regions of the dendrite (Fig. 1K). There was a trend for γ-tubulin-GFP accumulations to be located in more distal varicosities, but this is likely due to a higher frequency of distal varicosities. We also observed γ-tubulin-GFP within branchpoints (Fig. 1I,J), which have already been implicated in microtubule nucleation(Weiner et al., 2020; Thyagarajan et al., 2025). The accumulations of γ-tubulin appeared to be independent of membranous structures, such as endosomes or Golgi outposts as they tended to localise diffusely within varicosities and branchpoints (Fig 1I; Fig S1G). In contrast, the pH sensitive membrane reporter pHluorin-CD4-tdTom, previously used to track endosomes (Kanamori et al., 2015), appeared as discrete puncta that were not restricted to branchpoints and varicosities (Fig. S1G,H). Golgi outposts also localise as discrete puncta and are rare in these neurons (Mukherjee et al., 2020; Weiner et al., 2020).

In summary, the similar frequency and distribution of varicosities containing γ-tubulin and repeatedly initiating retrograde microtubule growth strongly suggest that a fraction of proximal, medial and distal dendritic varicosities are sites of microtubule nucleation that can generate minus-end-out microtubules.

### Microtubule minus ends accumulate specifically within distal varicosities

We next examined the localisation of minus ends within secondary dendrites using a modified form of a widely-used minus end reporter called Nod-LacZ (Clark et al., 1997). We remade this minus-end-directed motor fusion with fluorescent tags and confirmed the co-localisation of NodlacZ and Nod-sfGFP (Fig. S2A). Interestingly, Nod-sfGFP accumulated specifically within a selection of distal varicosities with varying patterns between dendrites (Fig. 2A,B). Multiple distal varicosities of an individual dendrites could be occupied by Nod-sfGFP (Fig S2B), suggesting that the motors don’t simply accumulate at the terminal, most distal, minus end within each dendrite. This localisation pattern was abolished when expressing a form mutated within its predicted microtubule binding region (UAS-Nod^6A^-sfGFP, see methods) (Fig. S2C). Consistent with an accumulation of minus ends specifically in distal varicosities, two MTOC proteins, Cnn and Ninein, both implicated in minus-end organization and anchoring, also accumulated specifically in distal varicosities (Fig. 2C-E, Fig. S2D-F), with Ninein colocalising perfectly with the minus end reporter (Fig. 2F).

**Figure 2.**
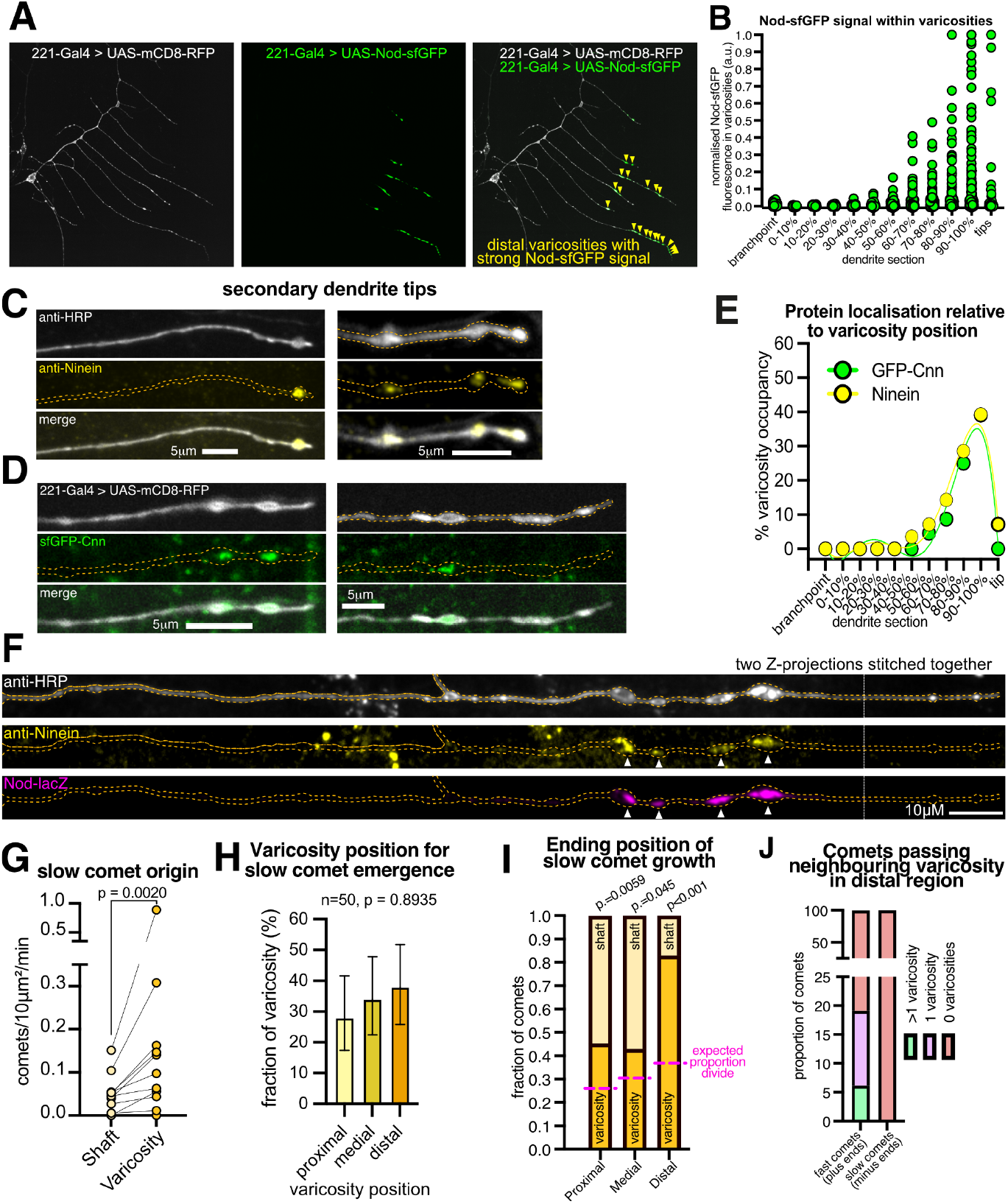
Microtubule minus ends accumulate in distal varicosities. (**A**) A confocal image of a dendritic arbor from a living 3^rd^ instar larval class I ddaE neurons expressing the membrane marker UAS-mCD8-RFP (grayscale) and the minus end marker UAS-Nod-sfGFP (green) with 221-Gal4. Note how Nod-sfGFP accumulates predominantly in distal varicosities. (**B**) Graph plotting the fluorescence intensity values of Nod-sfGFP within varicosities in relation to varicosity position. Each point represents a single varicosity. The data was normalised such that the highest value was equal to 1. N=1365 varicosities from 108 dendrites from 21 neurons. Note how the brightest intensities are found in distal varicosities, but that they are not confined to the most distal varicosities or dendrite tips. (**C**,**D**) Examples of distal secondary dendrites from a fixed (C) or living (D) 3^rd^ instar larval class I ddaE neuron either immunostained for HRP (grayscale) and Ninein (yellow) (C) or expressing the membrane marker UAS-mCD8-RFP (grayscale) with 221Gal4 and endogenously tagged sfGFP-Cnn (green) (D). (**E**) Example of a secondary dendrite from a fixed 3rd instar larval ddaE neuron immunostained for HRP (grayscale), Ninein (yellow), and lacZ (magenta). Arrowheads indicate the colocalisation of Ninein and Nod-lacZ within distal varicosities. White dotted line indicates the joining of two separate Z-projections. (**F**) Graph plotting the proportion of varicosities containing anti-Ninein signal (yellow, N=254 varicosities from 18 dendrites from 3 neurons) or sfGFP-Cnn signal (green, N=149 varicosities from 9 dendrites from 2 neurons) >20% above cytosolic background levels in relation to varicosity position. 5th order polynomial analyses generated trend lines are shown. Note that both Ninein and Cnn are enriched specifically in distal varicosities. (**G**) Graph displays the frequency at which slow EB1-GFP comets emerged from either shafts (light tone) or varicosities (dark tone). Each point represents the value from an individual dendrite. Lines join data from the same dendrite. Wilcoxon matched-pairs signed rank tests were used to compare the data from shafts and varicosities. (**H**) Graph displaying the spatial distribution of varicosities that have slow EB1-GFP comets emerging from them. A Chi^2^ test was performed to compare the observed proportions to what would be expected based on the total number of varicosities in each section. (**I**) Graph displays the proportion of slow EB1-GFP comets that stop growing within either shafts (light tone) or varicosities (dark tone). The magenta dashed lines indicate the expected proportions if minus ends were stopping randomly (these expected proportions are not at 50% due to difference in the proportion of the dendrite comprising shafts and varicosities). Chi^2^ tests were used to compare the observed and expected proportions. (**J**) Graph displays the proportion of either fast (left) or slow (right) EB1-GFP comets within distal regions that grow past neighbouring varicosities. Note that the Y-axis is split into two scales for better observation of the data. For G-J, data was obtained from 10 secondary dendrites from 8 neurons; 479 plus ends and 129 minus ends were recorded. Related to Figure S2.

We reasoned that minus ends may be enriched specifically in distal varicosities due to differences in minus end dynamics. Experiments *in vitro* have shown that minus ends can be released from γ-TuRCs, with either microtubule severing occurring close to γ-TuRCs (Henkin et al., 2023), or competition from CAMSAP/Patronin proteins (Rai et al., 2024). Consistent minus end release from γ-TuRCs within varicosities, we observed minus end growth initiating preferentially from varicosities compared to shafts (Fig. 2G). We did not, however, observe a preference for minus end release to occur from proximal and medial varicosities (Fig. 2H). Nevertheless, there was also a strong preference for minus ends to stop growing specifically within distal varicosities (Fig. 2I). Indeed, we never observed minus ends growing past a distal varicosity (Fig. 2J). This suggests that minus ends have a higher affinity for distal varicosities, consistent with the specific accumulation of the minus end reporter, Cnn and Ninein at these sites.

### Minus ends can accumulate in distal varicosities independently of Patronin-mediated minus end growth

It has been proposed that Patronin-mediated minus end growth is the mechanism that populates dendrites with minus-end-out microtubules in ddaE neurons (Feng et al., 2019). However, given that microtubule nucleation appears to occur at varicosities, minus-end-out microtubules could also be generated by distal nucleation sites. We therefore investigated whether eliminating minus end growth abolishes minus end accumulation in distal varicosities. Similar to primary dendrites (Feng et al., 2019), expressing Patronin-RNAi strongly reduced slow-moving minus end comets in secondary dendrites and increased the speed and frequency of fast-moving plus end comets (Fig. 3A-C); this speed increase in plus ends is presumably due to the induction of cellular stress pathways (Feng et al., 2019). Both primary and secondary dendrites also displayed polarity defects, with an increase in mixed or predominantly anterograde plus end growth (Fig. 3D). Nevertheless, dual-colour imaging of Nod-mCherry and EB1-GFP showed that Nod-mCherry could still strongly accumulate in the distal varicosities of correctly polarised dendrites in Patronin-RNAi neurons (Fig. 3E-G), despite slow moving comets not being observed in these dendrites. Nod-mCherry did not accumulate in the distal varicosities of incorrectly polarised dendrites from Patronin-RNAi neurons, or the few incorrectly polarised dendrites from control-RNAi neurons (Fig. 3F), but this is presumably because the minus-end directed motor is excluded from these dendrites. Thus, minus end growth is not essential for minus end accumulation in distal varicosities, so long as the dendrites are correctly polarised. This is consistent with minus-end-out microtubules being generated by local nucleation and anchoring events.

**Figure 3.**
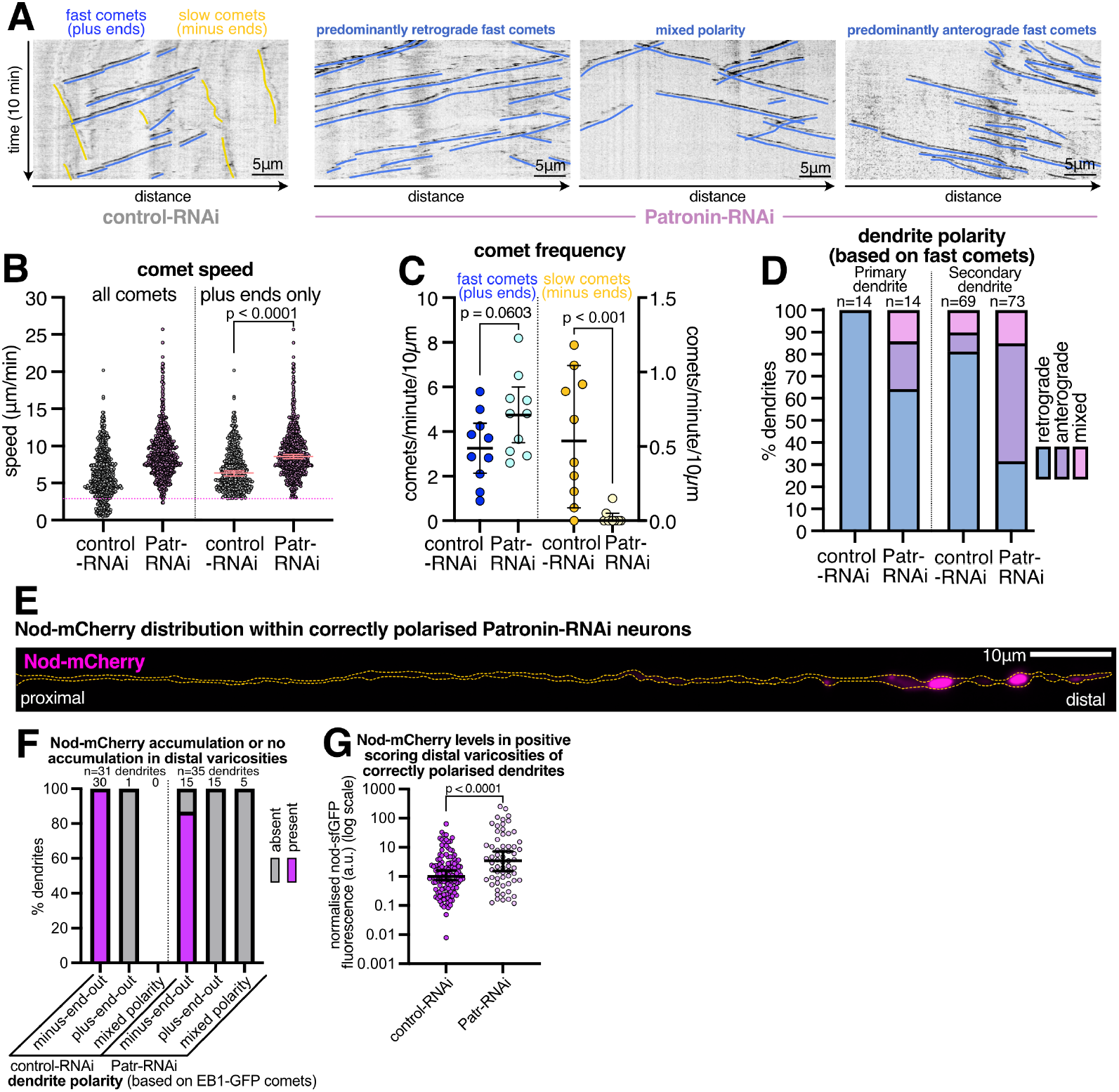
Patronin depletion prevents minus end growth and perturbs microtubule polarity, but does not prevent minus end accumulation in distal varicosities. (**A**) Example kymographs made from 10-minute videos of neurons from living 3rd instar larval class I ddaE neuron secondary dendrites expressing UAS-EB1-GFP and either a UAS-γ-tubulin37C-RNAi (control-RNAi) (left) or UAS-Patronin-RNAi (right 3 panels) with 221-Gal4. Different examples for Patronin-RNAi neurons are shown to highlight the variety of different polarities that the dendrites display. (**B**) Graph displaying the speed of individual EB1-GFP comets within secondary dendrites from either control-RNAi neurons or Patronin-RNAi neurons. Data on the right display only comets that have the speed cut off of 2.9 μm/min, i.e. plus ends. Medians with 95% CI are shown. A Mann-Whitney test was used to compare the speed of plus ends. (**C**) Graph comparing the frequency of fast-moving (left Y-axis) and slow-moving (right Y-axis) EB1-GFP comets in either control-RNAi or Patronin-RNAi neurons. Each point represents an individual dendrite. Means with 95% CIs are shown for fast comets; Medians with 95% CIs are shown for slow comets. An unpaired two-tail t-test was used to compare fast comet frequencies; a Mann-Whitney test was used to compare slow comet frequencies. For B,C control-RNAi data was obtained from analysing 513 EB1-GFP comets from 10 dendrites from 6 neurons; Patronin-RNAi data was obtained from analysing 616 EB1-GFP comets from 10 dendrites from 6 neurons. (**D**) Graph displaying the proportion of EB1-GFP comets (left) or secondary (right) dendrites that contained either predominantly retrograde or anterograde moving fast EB1-GFP comets or that had mixed polarity within proximal and medial regions, from control-RNAi and Patronin-RNAi neurons. N = 83 and 87 dendrites for control-RNAi and Patronin-RNAi, respectively, from 14 neurons. (**E**) Example images of a secondary dendrite that displayed predominantly minus-end-out microtubule polarity (based on EB1-GFP comet analysis) from a 3rd instar larval class I ddaE neuron expressing UAS-EB1-GFP and UAS-Nod-mCherry (magenta) with 221-Gal4. Note that Nod-mCherry accumulates strongly in distal varicosities, despite the absence of minus end growth. (**F**) Graph displaying the proportion of minus-end-out, plus-end-out, or mixed polarity secondary dendrites that contained accumulations of Nod-mCherry in distal varicosities from 3rd instar larval class I ddaE neurons expressing UAS-EB1-GFP, UAS-Nod-mCherry, and either UAS-control-RNAi (left) or UAS-Patronin-RNAi (right) with 221-Gal4. N for control = 31 dendrites from 10 neurons; N for Patronin-RNAi = 35 dendrites from 10 neurons. (**G**) Graph displaying the fluorescence intensity of the Nod-mCherry foci found within distal varicosities of correctly polarised control-RNAi or Patronin-RNAi neurons. Each dot represents a single varicosity. The data were normalised such that the median value for control-RNAi was equal to 1 and was plotted on a log scale. Medians and 95% CIs are shown. A lognormal t-test was used to compare the data.

### Ninein and Cnn are required for minus end organisation in distal varicosities

We hypothesised that Ninein and Cnn are important for minus end organisation in distal varicosities. Their individual depletion produced no clear phenotypes in the pattern of Nod-sfGFP (data not shown), but their co-depletion produced strong phenotypes, with microtubule minus ends forming striking buckles and loops within distal varicosities (Fig. 4A,B). This occurred within ∼30% of the varicosities that contained a significant Nod-sfGFP signal, far more frequently than in controls, Patronin-RNAi neurons, or in either single depletion condition (Fig. 4C). Varicosities containing microtubule loops were typically misshapen and enlarged, being approximately 5 to 6-fold larger in 2D projected area (Fig. 4B,D). Live imaging indicated that buckles and loops formed when microtubules pushed against the tips of dendrites, and this correlated with varicosity expansion (Fig. 4F,G; Videos S2,S3). Thus, compressive forces may lead to buckling and looping, similar to when microtubules push against the cell cortex in *Drosophila* wing epithelia (Singh et al., 2018). Similar microtubule loops have also been observed along axons after mis-regulation of growing plus ends (Hahn et al., 2021), perhaps due to a failure to properly organise microtubules into bundles (Voelzmann et al., 2016). In addition, dendrite length was more variable (Fig S3A), with dendrites containing microtubule defects being unusually long and those without defects unusually short (Fig. 4E).

**Figure 4.**
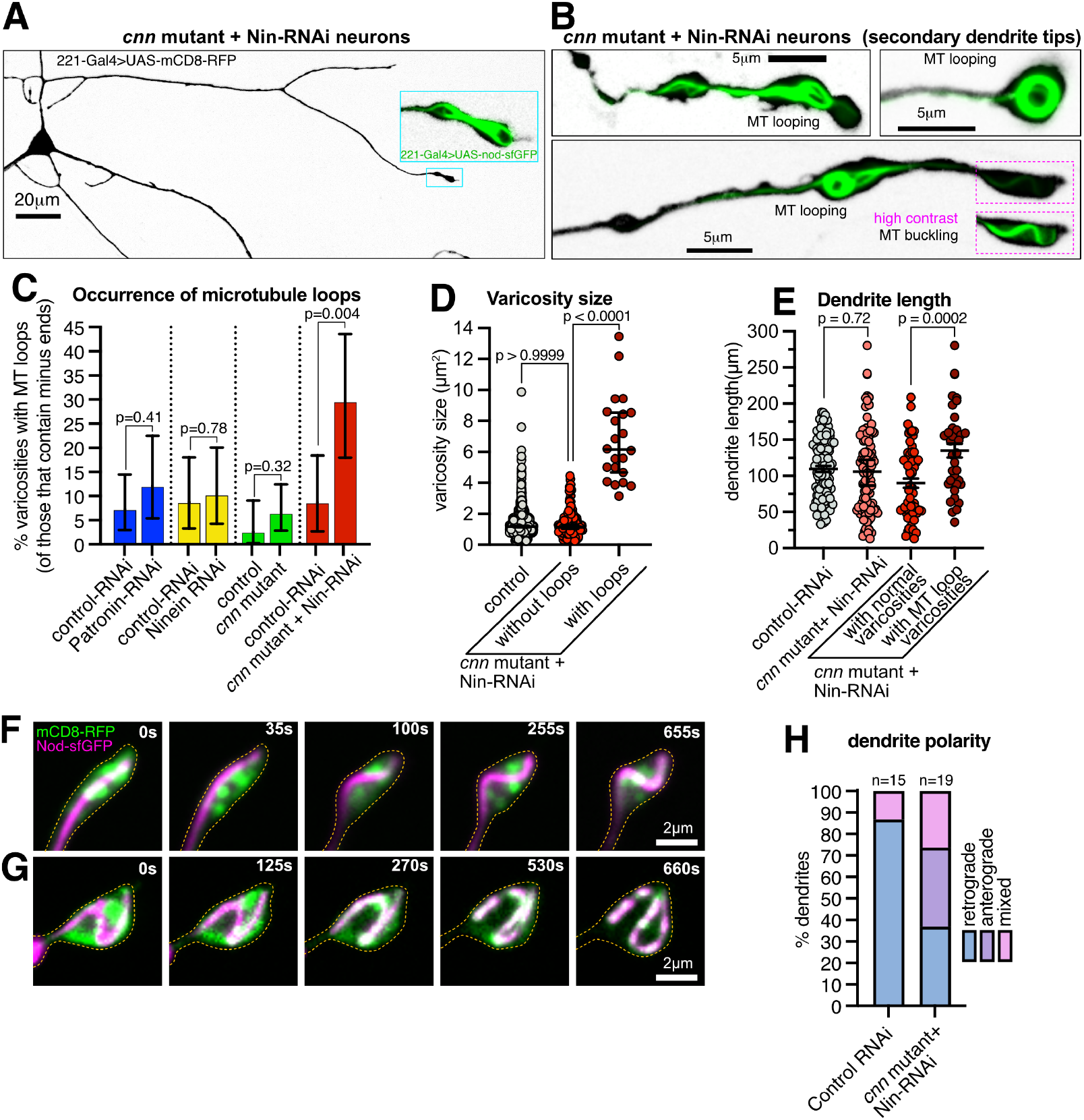
Ninein and Cnn are critical for minus end organisation in distal varicosities. **(A**,**B)** Confocal images of a secondary dendrite (A) and the distal regions of secondary dendrites (B) from class I ddaE neurons of *cnn* mutant 3rd instar larvae expressing UAS-Ninein-RNAi, UAS-mCD8-RFP (inverted grayscale) and UAS-Nod-sfGFP (green), i.e. Cnn/Ninein double-depletion neurons. Note how microtubule buckling and looping can be observed in distal varicosities. (**C**) Graph displaying the proportions of Nod-sfGFP-positive varicosities containing microtubule loops under various conditions, as indicated. N = 96, 66, 69, 68, 77, 123, 54 varicosities from 26, 22, 30, 20, 19, 24, 38 dendrites from 5, 5, 5, 5, 5, 5, 5 neurons for Control-RNAi (control for Patronin-RNAi), Patronin-RNAi, Control-RNAi (control for Ninein-RNAi), Ninein-RNAi, control (control for *cnn* mutant), *cnn* mutant, and *cnn* mutant + Ninein-RNAi neurons, respectively. Chi^2^ tests were used to compare the data. (**D**) Graph displaying varicosity size (measured as area of 2D image) in control and Ninein/Cnn double-depletion neurons, split by the presence (red) or absence of loops (dark red). Median and 95% CIs are shown. Each point represents a single varicosity. A Kruskal–Wallis test with multiple comparisons was used to compare the data. (**E**) Graph displaying secondary dendrite length in control or Cnn/Ninein double-depletion neurons, which is also split into two datasets: the presence (red) or absence of loops (dark red). Each point represents a single dendrite. Mean and SEM are shown. Mann–Whitney tests were used to compare the data. For D and E, N for control-RNAi neurons = 1365 varicosities from 108 dendrites from 21 neurons; N for Ninein/Cnn double-depletion neurons = 243 varicosities from 38 dendrites from 5 neurons. (**F**,**G**) Timelapse images from Video S2 (F) and Video S3 (G) of dendrite tips from neurons expressing UAS-mCD8-RFP (membrane; green) and UAS-Nod-sfGFP (magenta) with 221-Gal4. In (F), the minus end starts straight but then becomes buckled while contacting the distal dendrite tip; in (G), the minus end starts buckled and then starts to form a loop. (**H**) Graph displaying the proportion of secondary dendrites that contained either predominantly retrograde or anterograde growing plus ends or that had mixed polarity in proximal and distal regions from control and Cnn/Ninein double-depletion neurons. N = 15 dendrites from 6 neurons for control and 19 dendrites from 9 neurons for Cnn/Ninein double depletion. Related to Figure S3.

When expressing EB1-GFP instead of Nod-sfGFP, dendrites also displayed increased distal varicosity size compared to controls (Fig. S3B-D), and EB1-GFP comets took unusual paths within enlarged varicosities (Fig. S3C,D; Video S4, S5), consistent with microtubule defects occurring independent of the Nod-sfGFP reporter. There were also changes in microtubule growth, with an overall increase in EB1-GFP comet speed (Fig. S3E) and a reduction in the frequency of minus end growth (Fig. S3F). This latter observation suggested that excessive minus end growth was not the cause of microtubule buckling and looping and this was confirmed by expressing Patronin-RNAi in a Cnn/Ninein double depletion background and still observing a similar frequency of microtubule looping events (Fig. S3G). Furthermore, overall microtubule polarity was perturbed in Cnn/Ninein double depletion neurons, with only ∼37% of the secondary dendrites displaying minus-end-out polarity (Fig. 4H).

In conclusion, Ninein and Cnn are required to organise the minus ends of microtubules within distal varicosities to avoid microtubule disorganisation and dendrite morphology defects.

### A proposed model for microtubule generation and organisation in secondary dendrites of class I da neurons

A tentative model for the organisation of the microtubule network in *Drosophila* class I ddaE neuron secondary dendrites is shown in Fig. 5. This model is incomplete, but is consistent with our data.

**Figure 5.**
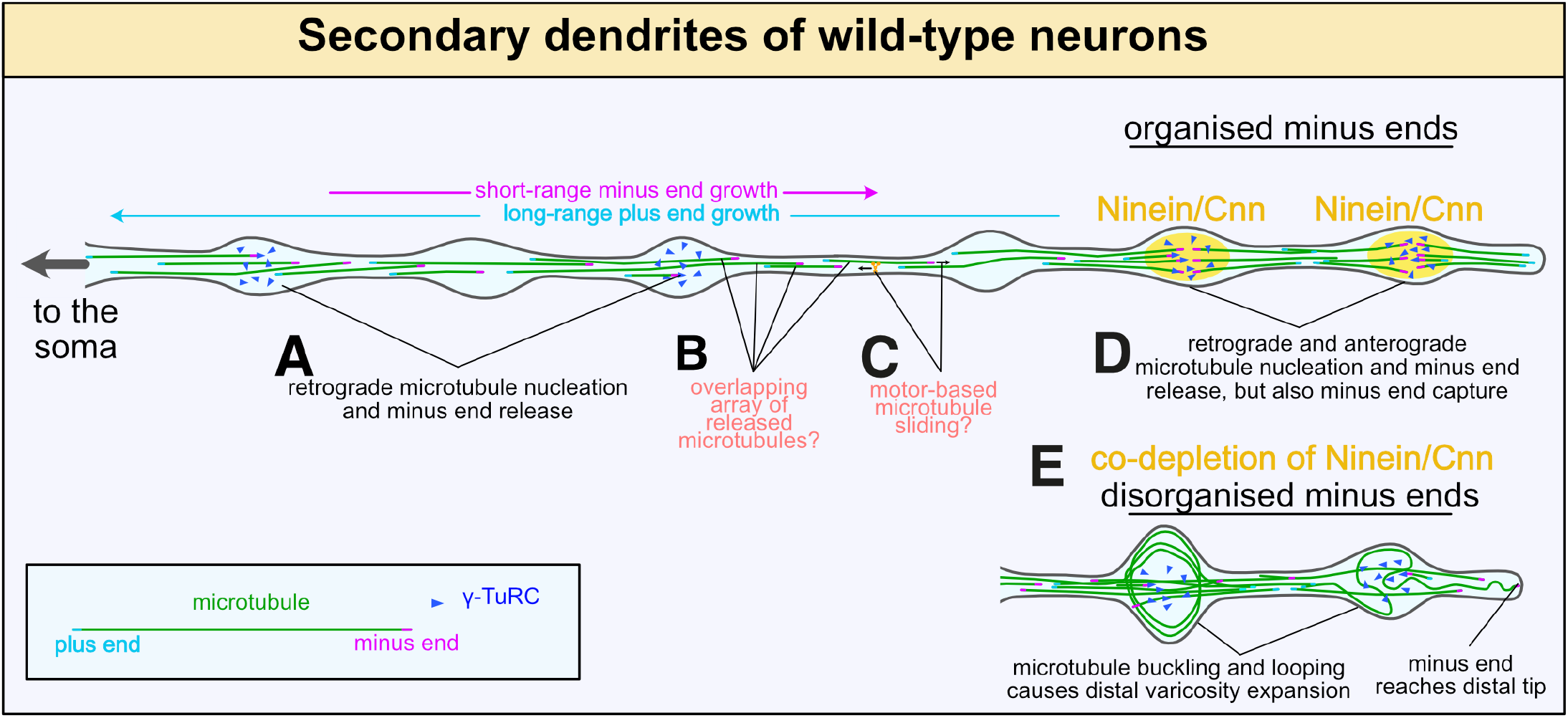
Model representing microtubule organisation in the secondary dendrites of ddaE neurons. This model represents our current working hypothesis based on the available data. While simplified and incomplete, it provides a mechanistic framework for microtubule generation and organization within these dendrites. γ-TuRCs (blue triangles) initiate microtubule nucleation from proximal and medial varicosities (A). Fast-growing plus ends grow long distances preferentially back towards the soma—the reason for the directional bias is unclear; minus ends are released from γ-TuRCs, but grow slowly and only short distances. The ultimate fate of these released microtubules is unclear and our data do not distinguish between different possibilities, e.g. depolymerisation; incorporation into a tiled overlapping array (B); or moved by molecular motors (C). Minus ends may also grow repeatedly to eventually reach distal regions, but our Patronin depletion analysis shows that this is not essential for minus-end accumulation in distal regions. γ-TuRCs also initiate microtubule nucleation from distal varicosities, but plus ends grow in both directions with equal frequency (D). Minus ends are released but they also preferentially stop growing and accumulate within distal varicosities, suggesting that they have a high affinity for these sites. This is consistent with the accumulation of two minus-end organising proteins, Cnn and Ninein, specifically in distal varicosities (D). Co-depletion of Cnn and Ninein leads to disorganisation of minus ends (E). Minus ends now reach the dendrite tip and appear to be pushed against it, leading to buckling and looping of the microtubule that distorts and expands varicosities (E). Why and how this occurs remains unclear, but it is not dependent on minus-end growth and we hypothesise it involves molecular motors that slide microtubules against the dendrite tip.

We propose that γ-TuRCs within proximal and medial varicosities nucleate microtubules whose plus ends preferentially grow back towards the soma, directly contributing to minus-end-out microtubule polarity. γ-TuRCs in distal varicosities also nucleate microtubules, but these grow bi-directionally. The reason for this difference remains unclear.

Minus ends appear to be released and grow short distances away from their nucleation sites, preventing their accumulation in proximal and distal varicosities. Based on *in vitro* experiments, minus end release from γ-TuRCs could occur via microtubule severing (Henkin et al., 2023) or competition with Patronin (Rai et al., 2024). The fate of these released minus ends is unclear – they could be depolymerised, regrow or the microtubules could be transported by motors such that the minus ends reach distal varicosities.

In contrast to more proximal varicosities, minus ends accumulate and are organised in distal varicosities via the specific accumulation of two MTOC and minus end organising proteins, Cnn and Ninein. Both proteins can associate with γ-TuRCs (Tovey et al., 2021; Zheng et al., 2016) and Ninein can bind microtubules (Kowanda et al., 2016). Moreover, Cnn and its homologues can modify γ-TuRC activity, both increasing their nucleation potential (Choi et al., 2010; Tovey et al., 2021; Serna et al., 2024; Rale et al., 2022; Xu et al., 2024) and inhibiting minus end release by Patronin family members (Rai et al., 2024). Cnn can also form scaffolds that recruit other MTOC proteins (Conduit et al., 2014a; b), which could in turn promote minus end retention. Cnn and Ninein are therefore well placed to organise minus ends in distal varicosities. While the precise molecular mechanisms underlying their involvement remain to be determined, it is very clear they are required to prevent minus ends disorganising into buckles and loops that distort the neuronal membrane. The forces that induce this buckling and looping are unclear, but microtubule sliding has already been reported in *Drosophila* neurons (Lu et al., 2013) and could certainly be involved.

The accumulation of minus ends in distal varicosities does not require minus end growth, and so we expect local nucleation and anchoring are the main driving forces behind this phenomenon. Nevertheless, eliminating minus end growth via Patronin depletion does lead to a higher fraction of mis-polarised dendrites, similar to the effect of co-depleting of Cnn and Ninein. Thus, both minus end growth and minus end anchoring in distal varicosities may collectively help establish microtubule polarity during early dendrite development.

In summary, we identify dendritic varicosities as a new neuronal nc-MTOC and demonstrate their importance for regulating the polarised dendritic microtubule network in the secondary dendrites of Drosophila ddaE neurons.

## Acknowledgements

This research was supported by the Centre National de la Recherche Scientifique (CNRS), by a Chaire d’excellence grant from the IdEx Université Paris Cité (ANR-18-IDEX-0001), by an ATIP Avenir award funded by the Fondation Bettencourt Schueller, by a “Equipe FRM 2023” grant (EQU202303016314), by an Impulscience grant funded by the Fondation Bettencourt Schueller, and by a PhD fellowship awarded by the Bio-SPC école doctorale, Université Paris Cité. We thank Jordan Raff for kindly donating antibodies. We thank Arthur Molines for providing the macro for the gaussian blur denoising filter. We thank members of the Conduit lab for their critical reading of the manuscript and intellectual input. The work benefited from the ImagoSeine at the IJM, Paris. This paper was typeset with the bioRxiv word template by @Chrelli: www.github.com/chrelli/bio-Rxiv-word-template. For the purpose of Open Access, a CC-BY public copyright license has been applied by the authors to the present document and will be applied to all subsequent versions up to the Author Accepted Manuscript arising from this submission.

## Author contributions

PTC conceived and supervised the project and obtained funding. AC and CV started and finished the project, respectively. AC analysed varicosity distribution, performed heat fixation, generated the fluorescent Nod lines, determined the distribution of proteins in varicosities, and documented the phenotypes in Cnn/Ninein double depleted neurons expressing Nod-sfGFP. CV performed all the microtubule dynamics experiments and analysed the data across all conditions and performed and analysed all the experiments for Patronin-depleted neurons. CV performed some reanalysis of data generated by AC. PTC analysed the data for the distribution of pHluorin-CD4-tdTom puncta. PTC and CV wrote the manuscript.

## Competing interest statement

The authors declare no competing interests.

## Materials and Methods

### Drosophila handling and stocks

All *Drosophila melanogaster* (RRID:NCBITaxon_7227) strains were maintained at 18 or 25◦C on Iberian fly food made from dry active yeast, agar, and organic pasta flour, supplemented with nipagin, propionic acid, pen/strep and food colouring.

The following fluorescent alleles were used in this study: γ-tubulin23C-sfGFP(Tovey et al., 2018), γ-tubulin23c-eGFP(Mukherjee et al., 2020), UAS-EB1-GFP (BL 35512), sfGFP-CnnP1, UAS-mCD8-RFP (BDSC 27392), UAS-Nod-sfGFP (this study), UAS-Nod-mCherry (this study), and pUASt-Nod^6A^-sfGFP (this study). The following mutant alleles were used in this study: cnn^f04547^, which prevents expression of the long Cnn-P1 isoform. The following Gal4, mutant, RNAi and Dicer lines were used in this study: 221-Gal4 (BL 26259) and 109-Gal4 (BDSC 8769). For control-RNAi, either γ-tubulin37C-RNAi (BL32513) or UAS-Rtnl2-RNAi (BL 58208) was used. For RNAi of specific genes we used: Patronin-RNAi (BL 36659), UAS-Ninein-RNAi (VDRC 13998).

For RNAi experiments, a single copy of the UAS-RNAi construct was expressed using 221-Gal4. For fluorescent protein localisation experiments, one copy of UAS-mCD8RFP was expressed to highlight cell membranes. For examining the localisation of γ-Tubulin23C, flies expressed one copy of γ-Tubulin23C-sfGFP together with one copy of γ-Tubulin23C-ssss-eGFP. For examining the localisation of Centrosomin-P1, two copies of endogenously tagged sfGFP-Cnn-P1 were expressed. For examining minus-end localisation, one copy of UAS-Nod-lacZ, UAS-Nod-sfGFP, UAS-Nod-mCherry was expressed with 221-Gal4. For analysing microtubule dynamics, flies expressed one copy of UAS-EB1-GFP with 221-Gal4.

### Generation of Transgenic Lines

For generating the pUASt-Nod-sfGFP and pUASt-Nod-mCherry transgenic lines, we first created a Gateway entry vector containing the Nod motor domain fused to Kinesin heavy chain (made by gene synthesis, ThermoFisher) using the same sequence as used in Clark et al., 1997 for Nod-LacZ. We modified a UASt Gateway destination vector that contained an attB genome integration site (gift from Matthieu Sanial) by introducing either the sfGFP or mCherry coding sequence via PCR and HiFi DNA Assembly (NEB). The final expression vectors containing the nod-kinesin coding sequence tagged with sfGFP or mCherry were generated via Gateway cloning (ThermoFisher). The vectors were amplified using a Midiprep kit (Qiagen) and sent to BestGene, Inc for injection into lines containing an appropriate attB “landing site” (2^nd^ Chromosome, attp40; 3^rd^ Chromosome, attP2) allowing integration via the PhiC31 integrase system.

The pUASt-Nod^6A^-sfGFP line was generated by introducing the following mutations into the Nod motor domain: K148A, R165A, R168A, R234A, R243A, R273A. These residues were chosen based on the findings from Woehlke et al.(Woehlke et al., 1997) about how Kinesin1’s head domain interacts with microtubules and an overlay of AlphaFold2 predictions of the Nod and Kinesin1 motor heads using colabfold (Mirdita et al., 2022) in ChimeraX (Meng et al., 2023).

### Immunostaining

Dissected larvae were processed as previously described(Mukherjee et al., 2020). Briefly, fillet preparations were fixed in freshly prepared 4% formaldehyde for 20 min at room temperature and were then washed four times for 10 min in PBS. Blocking was carried out in PBS-BT (PBS plus 0.2% Triton X-100 (PBST) and 5% BSA) for 1h at room temperature. Preparations were incubated with appropriate primary antibodies diluted in PBS-BT overnight at 4ºC. After washing in PBST for ∼8h, changing washes every 30-45 min, samples were incubated in secondary antibodies diluted in PBS-BT overnight at 4ºC. The fillet preparations were then washed for ∼8h, changing washes every 30-45 min in PBST before mounting in Mowiol. They were directly stored at -20ºC and imaged within a week. Neurons within segments A2 and A6 were imaged.

### Heat fixation

Larvae were placed in a drop of glycerol on top of a glass slide and placed on a 70°C heat block for 10 seconds. The anterior end of the larvae was then removed and the larvae was gently squashed between the slide and a coverslip before imaging.

### Antibodies

The following primary antibodies were used: anti-GFP mouse monoclonal at 1:250 (Roche, 11814460001), anti-Ninein guinea pig at 1:500 (gift from the Lecuyer Lab), Alexa-647 HRP conjugated polyclonal at 1:500 (Jackson), and anti-*β*-galactosidase rabbit polyclonal at 1:2500 (Invitrogen A11132). The following secondary antibodies were used: Alexa-488 anti-mouse at 1:500 (ThermoFisher), Alexa-547 anti-rabbit at 1:500 (ThermoFisher), and Alexa-488 anti-guinea pig (ThermoFisher).

### Microscopy

All imaging was carried out at ambient temperature (∼21ºC). For live larval samples, wandering third instar larvae were first washed in PBS and then placed in a drop of glycerol before being flattened between a slide and a 22X22mm coverslip, held in place by tape, and then imaged immediately for a maximum of 5 min. For observing microtubule dynamics using EB1-GFP, imaging was carried out on a Nikon Ti2 inverted microscope associated with a CSU-W1 T1 Yokogowa spinning disk system with a Kinetix 22 camera (Photometrics) run by Gataca’s lightshell for µManager software using a 60×1.4NA oil immersion lens (Nikon). Z-stacks with 1µm spacing were acquired every 3 seconds. Fixed and live imaging for protein localisation was carried out on one of 3 microscopes: a Zeiss Axio Observer.Z1 inverted microscope associated with a CSU-X1 Yokogowa spinning disk system with an ORCA Fusion camera (Hamamatsu) run by Zeiss ZenBlue acquisition software using a 60×1.4NA oil immersion lens (Zeiss); a Nikon Ti2 inverted microscope associated with a CSU-W1 T1 Yokogowa spinning disk system with a Kinetix 22 camera (Photometrics) run by Gataca’s lightshell for Micromanager software using a 60×1.4NA oil immersion lens (Nikon); or a Olympus FV3000 scanning inverted confocal system run by FV-OSR software using a 60X 1.4NA silicon immersion lens (UPLSAPO60xSilcon). Z-stacks with 0.5µm spacing were acquired. All images were processed in Fiji (Image J).

### Image Analysis

Fluorescent values, distances and areas were obtained in Fiji using the provided tools. The data was subsequently processed in Microsoft Excel, but statistical analysis and graph production were performed in GraphPad Prism. Datasets were assessed for Normality and lognormality before deciding whether to use a parametric or non-parametric test. A correction for multiple comparisons was used when necessary. Details of specific tests used are given in the Fig. legends. When determining varicosity position, if a secondary dendrite branched into two, we measured the position of the sections within each branch relative to the primary branchpoint and not from the secondary branchpoint.

During EB1-GFP comet analysis to examine microtubule nucleation, comets that were present within the first timepoint were excluded. Kymographs were generated using the plugin in Fiji. When producing Videos 1, 4 and 5, a Gaussian blur denoising filter was applied in which the intensity value of each pixel was replaced with a weighted average of its neighbouring 3 pixels.

For analysing the intensity of fluorescent signals within varicosities or branchpoints, maximum intensity Z-plane projections were first made and sum fluorescence within the appropriate ROI was obtained. A mean cytosolic signal was obtained from proximal regions of each dendrite and was used to perform a background subtraction such that any branchpoint or varicosity that had a similar cytosolic signal would have a value of ∼0. The data was then normalised such that the brightest signal had a value of 1. When quantifying the occupancy of proteins within branchpoints and varicosities, a positive occupancy was scored when the fluorescence signal within the structure was at least 20% higher than the cytoplasmic baseline.

**Figure S1.**
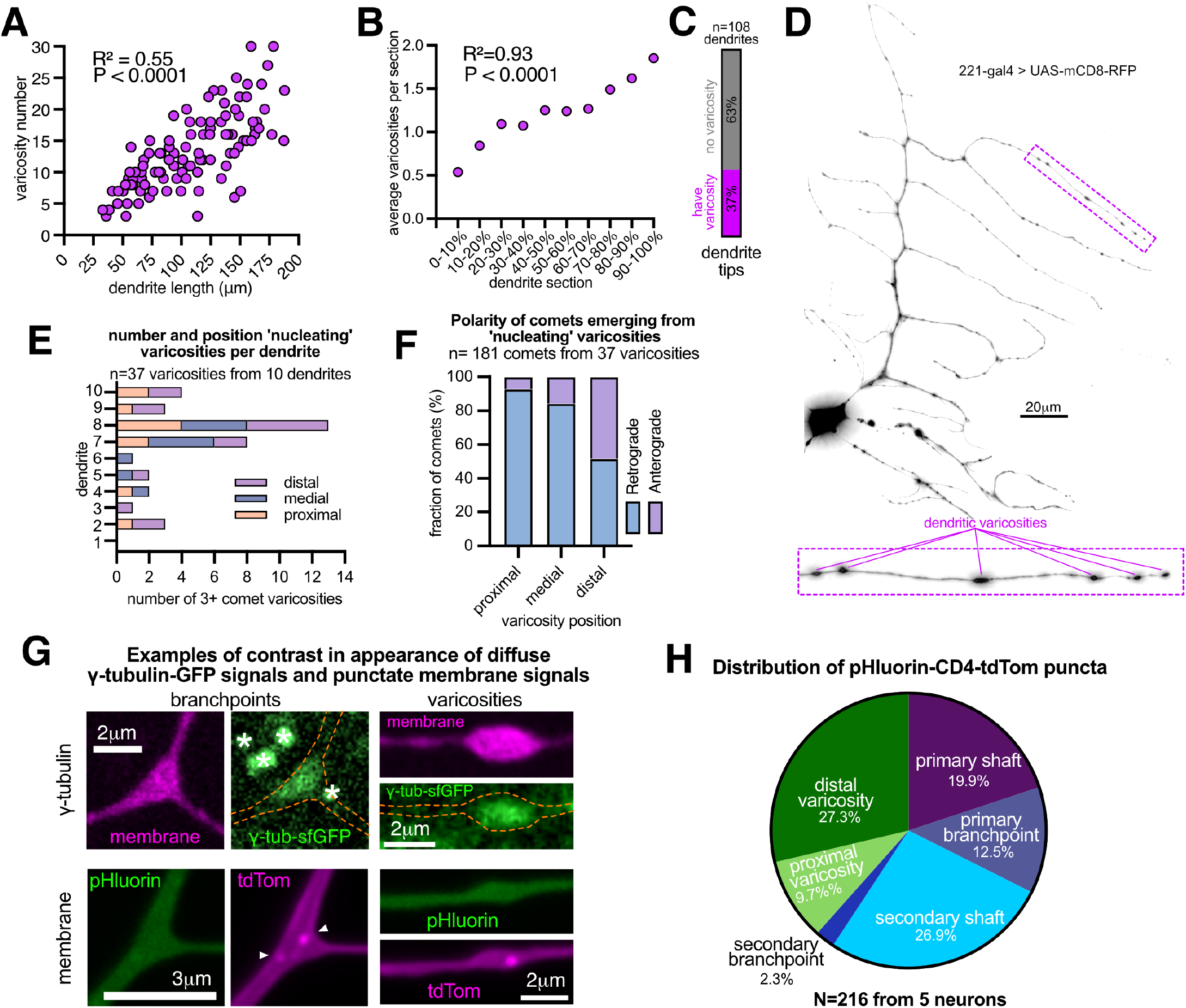
Characterisation of dendritic varicosities. **(A)** Graph plotting varicosity number versus dendrite length. R^2^ and P values were obtained by simple linear regression analysis. The positive linear correlation suggests that as dendrites grow more varicosities are added. (**B**) Graph plotting the average number of varicosities observed in relation to varicosity position (represented by 10% dendrite length sections, with 0% being the branchpoint). R^2^ and P values were obtained by simple linear regression analysis. The positive linear correlation shows that varicosity density increases along the dendrite length. (**C**) Graph displaying the proportion of dendrite tips that display a varicosity. In A–C, N = 1365 varicosities from 108 dendrites from 21 neurons. (**D**) A confocal image of a class I ddaE neuron dendritic arbor and an enlarged portion of a secondary dendrite from a rapidly heat-fixed 3rd instar larva expressing the membrane marker UAS-mCD8-RFP (inverted grayscale) with 221-Gal4. Note how dendritic varicosities can be observed along secondary dendrites, confirming they are observed under multiple conditions (chemical fixation, heat fixation and live imaging). (**E**) Graph displaying the position of microtubule nucleating varicosities within individual dendrites. n = 37 varicosities from 10 dendrites from 8 neurons. (**F**) Graph displays the polarity of fast EB1-GFP comets emerging from “nucleating” varicosities. N = 181 comets from 37 varicosities. (**G**) Confocal images of primary branchpoints and varicosities from 3rd instar class I dda neurons expressing either endogenously tagged γ-tubulin-sfGFP (green) and UAS-mCD8-RFP (magenta), or the pH-sensitive membrane reporter UAS-pHluorin-CD4-tdTom with 221-Gal4 (green = fluorescence lost at low pH, such as within late endosomes; magenta is tdTom fluorescence). Asterisks mark additional γ-tubulin-GFP signal from outside the neuron. Note that γ-tubulin-sfGFP within the branchpoint and varicosity is diffuse, while the membrane vesicle fluorescence is punctate. (**H**) A pie chart displaying the proportion of red-only pHluorin-CD4-tdTom foci in different regions of the dendrite, as indicated. Note that ∼47% of foci are located within shafts, where we never observe accumulations of γ-tubulin-GFP. Related to Figure 1.

**Figure S2.**
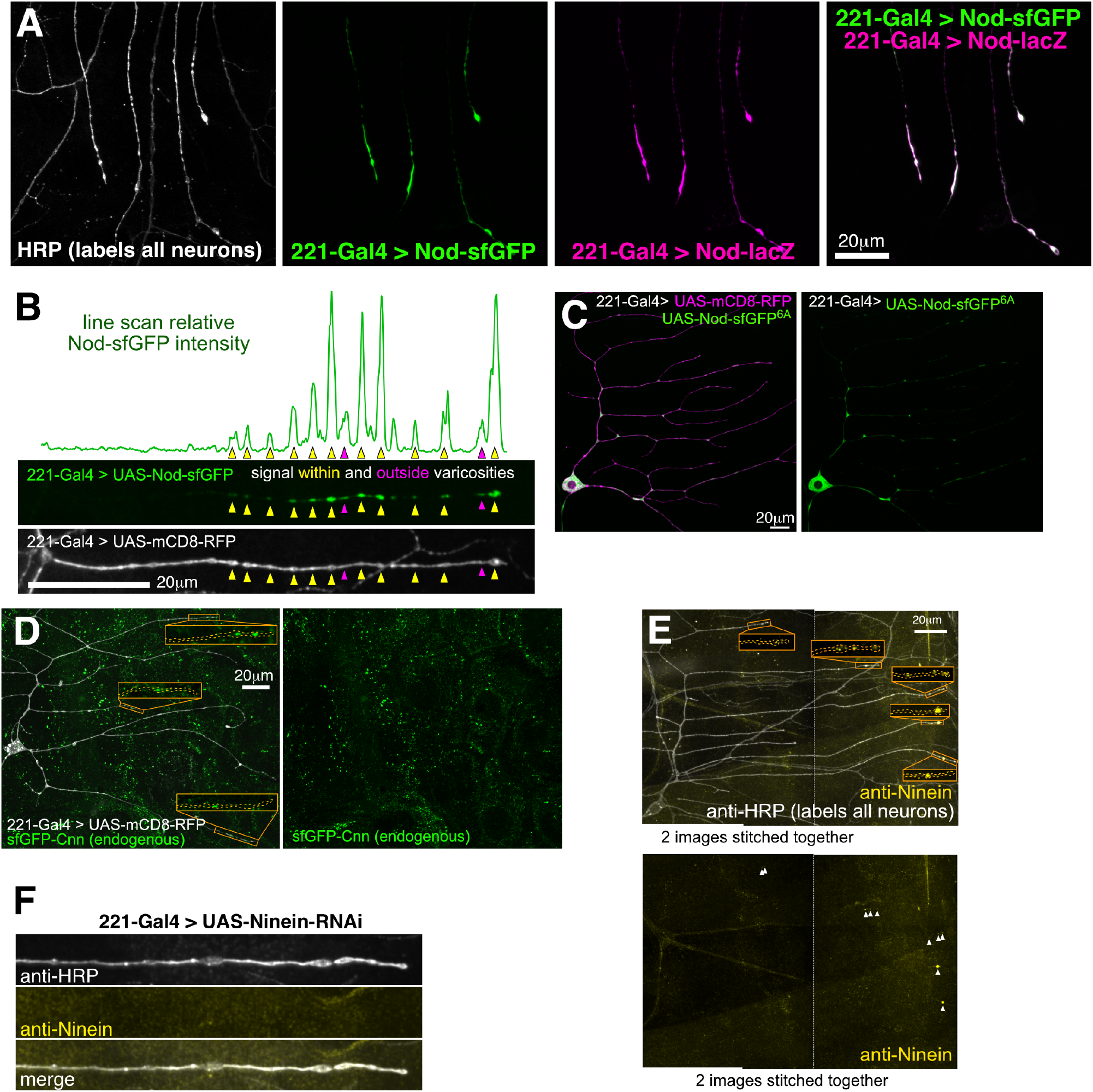
The minus end reporters Nod-sfGFP and Nod-lacZ and the MTOC proteins Ninein and Cnn localise specifically within distal varicosities. **(A)** Confocal images of secondary dendrites from a fixed 3rd instar larval ddaE neuron co-expressing UAS-Nod-sfGFP (green) and UAS-Nod-lacZ (magenta) with 221-Gal4, immunostained for HRP (neuronal membranes) and lacZ. Nod-sfGFP and Nod-lacZ colocalise. **(B)** Example of a secondary dendrite from a living 3rd instar larval class I ddaE neuron expressing the membrane reporter UAS-mCD8-RFP (grayscale) and the minus-end reporter UAS-Nod-sfGFP (green) with 221-Gal4. The associated fluorescence line plot shows that the strongest Nod-sfGFP signals are within distal varicosities (yellow arrowheads). Weaker, but still significant, signal can sometimes be observed outside varicosities (magenta arrowheads). **(C)** Confocal images of a dendritic arbor from a living 3rd instar larval ddaE neuron expressing UAS-mCD8-RFP (magenta) and a mutant form of the minus-end reporter UAS-Nod^6A-sfGFP (green) with 221-Gal4. The six alanine mutations are predicted to abolish microtubule binding. The mutant protein accumulates predominantly in the soma and also floods branchpoints, typical of overexpressed proteins in these neurons, but does not accumulate in distal varicosities. **(D, E)** Confocal images of dendritic arbors from either living (D) or fixed (E) 3rd instar larval ddaE neurons either expressing endogenously tagged sfGFP-Cnn (green) and UAS-mCD8-RFP (grayscale) with 221-Gal4 (D) or immunostained for Ninein (yellow) and HRP (grayscale) (E). Overlays are shown on the left and the individual sfGFP-Cnn (D) and anti-Ninein (E) channels are shown on the right. Orange boxes are zoomed in and re-contrasted images of distal dendrite regions to highlight the accumulation of sfGFP-Cnn and Ninein in distal varicosities. Arrowheads indicate Ninein accumulations in (E). Note that sfGFP-Cnn puncta are present in surrounding epidermal cells. **(F)** Confocal images of a secondary dendrite from a fixed 3rd instar larval ddaE neuron immunostained for HRP (grayscale) and Ninein (yellow) expressing UAS-Ninein-RNAi with 221-Gal4, showing the absence of a Ninein signal. Related to Figure 2.

**Figure S3.**
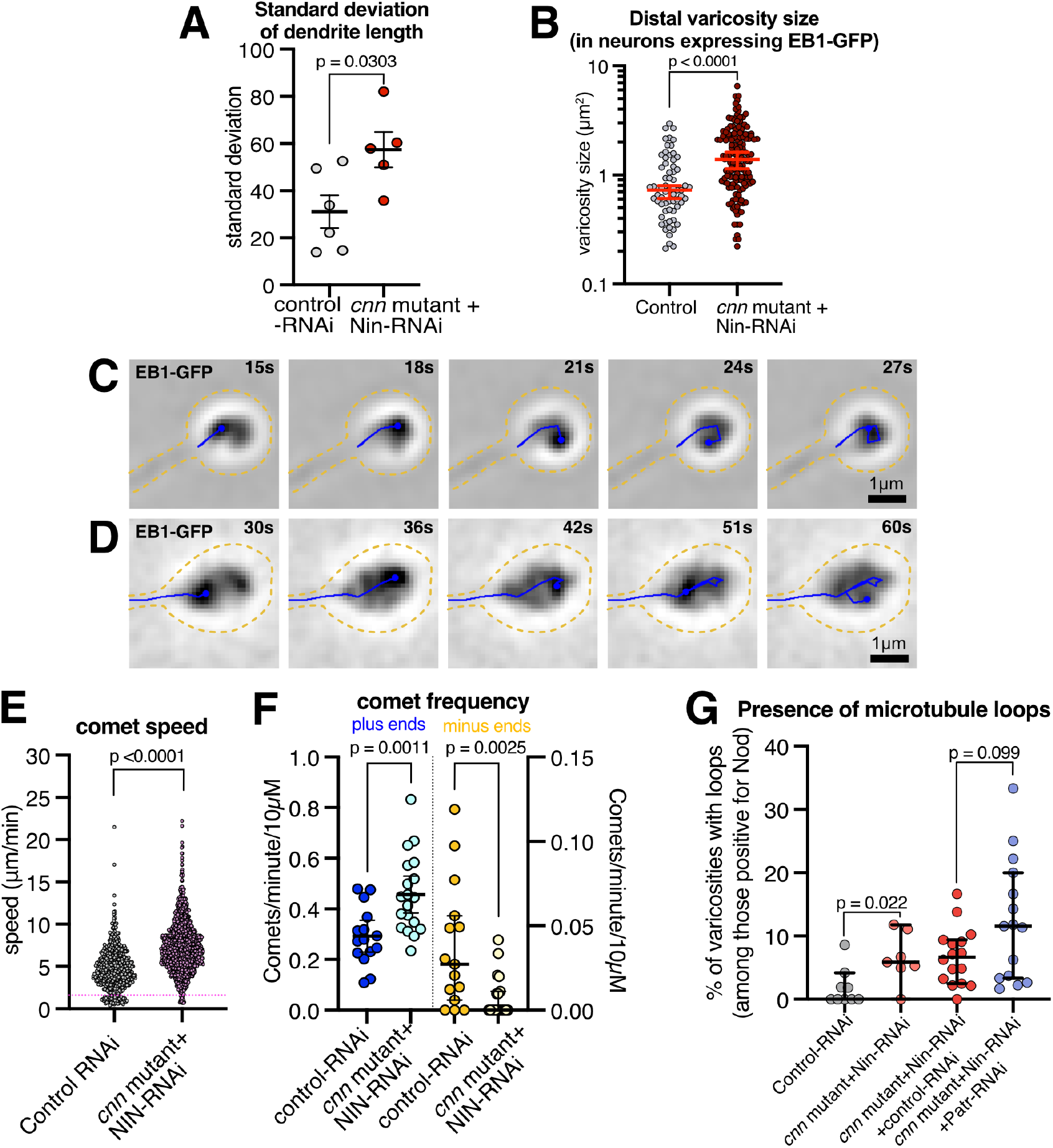
Phenotypes in Cnn/Ninein double-depletion neurons and how they are not rescued by eliminating minus end growth. **(A)** Graph displaying the standard deviation of dendrite length (i.e. how variable dendrite length is) in either control (grey) or Cnn/Ninein double-depletion neurons (red). Mean and SEM are shown. **N:** 55 or 38 dendrite tips from 6 or 5 neurons for control-RNAi and *cnn* mutant + Ninein-RNAi neurons, respectively. **(B)** Graph displaying distal varicosity size in either control or Cnn/Ninein double-depleted neurons expressing EB1-GFP instead of Nod-sfGFP. Unlike Figure 4D,E, the data could not be split into varicosities that did or did not contain microtubule loops. The data are lognormally distributed and plotted on a log scale. Medians with 95% confidence intervals are shown. A Welch’s *t*-test was used to compare the data. **N:** 66 varicosities from 19 secondary dendrites in 7 neurons from the control and 151 varicosities from 43 secondary dendrites in 12 neurons from the Cnn/Ninein double-depleted condition. **(C, D)** Timelapse images from Video S4 (C) and Video S5 (D) of dendrite tips from Cnn/Ninein double-depleted neurons expressing UAS-EB1-GFP (inverted grayscale) with 221-Gal4. The images were processed with a Gaussian blur filter to increase the signal-to-noise ratio. Blue lines indicate the paths of EB1-GFP comets that take unusual “looping” paths within the varicosities. **(E)** Graph displaying the speed of individual EB1-GFP comets within secondary dendrites from either control or Cnn/Ninein double-depletion neurons. The dotted line in magenta indicates the speed cut-off between the two populations of fast and slow comets. A Mann–Whitney test was used to compare the data. **N:** For (E), control-RNAi data were obtained from analysing 487 EB1-GFP comets from 15 dendrites from 6 neurons; Cnn/Ninein double-depletion data were obtained from analysing 1132 EB1-GFP comets from 19 dendrites from 9 neurons. **(F)** Graph comparing the frequency of growing plus (left Y-axis) and minus (right Y-axis) ends in either control or Cnn/Ninein double-depletion neurons. Each point represents an individual dendrite. Means with 95% confidence intervals are shown for plus ends; medians with 95% confidence intervals are shown for minus ends. An unpaired two-tail *t*-test was used to compare plus-end frequencies; a Mann–Whitney test was used to compare minus-end frequencies. **(G)** Graph displaying the proportion of varicosities containing a positive Nod-sfGFP signal that contained microtubule loops in different conditions, as indicated. The *cnn* mutant + Ninein-RNAi + control-RNAi condition was included to ensure balance between the number of UAS constructs when comparing to the *cnn* mutant + Ninein-RNAi + Patronin-RNAi condition. Each point represents an individual neuron. Medians with 95% confidence intervals are shown. Mann–Whitney tests were used to compare the data. Note how the expression of UAS-Patronin-RNAi, which eliminates minus-end growth, does not prevent the formation of microtubule loops in Cnn/Ninein double-depletion neurons. N: 11, 7, 16, 15 for control-RNAi, *cnn* mutant + Ninein-RNAi, *cnn* mutant + Ninein-RNAi + control-RNAi, and *cnn* mutant + Ninein-RNAi + Patronin-RNAi, respectively. Related to Figure 4.

## Notes

### Competing Interest Statement

The authors have declared no competing interest.

